# LINKING MULTI-SCALE BRAIN CONNECTIVITY WITH VIGILANCE, WORKING MEMORY, AND BEHAVIOR IN ADOLESCENTS

**DOI:** 10.1101/2025.05.09.653144

**Authors:** Prerana Bajracharya, Ram Ballem, Jiayu Chen, Pablo Andrés-Camazón, Nigar Khasayeva, Vince Calhoun, Jingyu Liu, Armin Iraji

## Abstract

This study examines how multi-scale intrinsic connectivity networks (ICNs) relate to cognitive and behavioral functions in adolescents, focusing on attention/vigilance, working memory, and behavioral regulation. Leveraging the NeuroMark 2.2 multi-scale ICN template obtained from over 100,000 subjects, we obtained multi-scale ICNs from baseline resting-state fMRI data from the ABCD Study. For this study, we are interested in “the fronto-thalamo-cerebellar (FTC) circuitry” and choose the subdomains of Neuromark 2.2 that cover it: Cerebellar (CB), Subcortical - Extended Thalamic (SC-ET), Higher Cognition - Insular Temporal (HC-IT), and Higher Cognition - Frontal (HC-FR), previously identified as relevant to cognitive and behavioral functions. Employing a multivariate approach combining principal component analysis (PCA) and canonical correlation analysis (CCA), we examined associations between these multi-scale ICNs and cognitive-behavioral outcomes. Our findings revealed significant associations, particularly for one of the estimated canonical components, linking multi-scale ICNs to cognitive and behavioral measures across both discovery and replication sets. This connectivity pattern may serve as a potential marker for attention, working memory, and behavioral regulation, offering new insights into a wide spectrum of neurodevelopmental disorders including Attention-Deficit/Hyperactivity Disorder (ADHD).

## 1. INTRODUCTION

ADHD and other neurodevelopmental disorders exhibit complex cognitive impairments in attention/vigilance, working memory, and behavioral regulation [1]. These cognitive functions are linked to interactions within and between multiple ICNs in the brain, such as the frontoparietal network, dorsal attention network, and default mode network [2], [3]. However, it remains unclear how ICNs at various functional scales contribute to cognitive variability and behavioral outcomes. Additionally, the association with fronto-thalamo-cerebellar (FTC) circuitry is sparsely researched. Understanding the neural correlation may yield insights into the brain’s functional organization and mechanisms underlying cognitive dysfunction.

Resting-state functional magnetic resonance imaging (rsfMRI), enables multi-scale analysis of ICNs, revealing their role in cognitive processes [4], [5]. Studying ICNs at different spatial resolutions—ranging from large [2], global networks to more localized subnetworks—has demonstrated associations with cognitive capacities, suggesting that brain organization is intrinsically tied to attentional and working memory processes. The 0-back task in the n-back paradigm has been widely used to assess vigilance by measuring basic attention, while the 2-back condition provides a measure of working memory capacity [6], [7]. Additionally, the Child Behavior Checklist (CBCL) allows for a comprehensive assessment of externalizing and internalizing behaviors, which reflect broader aspects of cognitive and emotional functioning in adolescents [8], [9]. Investigating ICN connectivity across spatial scales may clarify their contributions to cognition and behavior.[8].

The NeuroMark 2.2 template, derived from rsfMRI data of over 100,000 individuals [10], offers a robust representation of multi-scale ICNs, categorizing 105 ICNs into 7 domains and 14 subdomains [11]. This study focuses on four subdomains— cerebellar (CB), subcortical extended thalamic (SC-ET), higher cognition insular temporal (HC-IT), and higher cognition frontal (HC-FR)—which encompass the FTC circuitry, an area linked to cognitive and behavioral regulation. This focus was informed by prior research [12], [13], that underscored the importance of cerebellar-subcortical-cortical pathways in cognitive regulation and provides insights into neuroanatomical bases for cognitive processing.

In this study, we leverage a principal component analysis (PCA) + canonical correlation analysis (CCA) how static functional network connectivity (sFNC) between multi-scale ICNs relates to cognitive and behavioral measures. PCA+CCA is a multivariate technique that reduces dimensionality and reveals latent relationships among high-dimensional data. The sFNC captures the temporal interdependence among ICN time courses [14]. By integrating sFNC with neurocognitive and behavioral measures using PCA+CCA, this study aims to elucidate the neurobiological bases of attention/vigilance, working memory, and CBCL-reported behavior. We hypothesize that unique ICN configurations at different scales will correlate with distinct cognitive and behavioral profiles, revealing brain-cognition relationships fundamental to understanding both typical and atypical neurodevelopment.

This research aims to advance the understanding of brain-behavior relationships in adolescents by combining multi-scale network analysis with statistical methods to reveal latent factors underpinning cognitive functions and behavioral outcomes. This approach holds the potential to study ICN contributions to attention and working memory and to provide markers for neurodevelopmental disorders characterized by deficits in these domains.

## 2. METHODS AND MATERIALS

### 2.1. Dataset

This study utilized data from the ABCD Study (release 4.0), a population-based sample of 9-to 10-year-olds from 21 U.S. sites, including neuroimaging and behavioral data collected at baseline and longitudinally [8]. After rigorous quality control, 7,655 participants were included.

### 2.2. Preprocessing

We preprocessed the raw resting-state fMRI data using the FMRIB Software Library (FSL) v6.0 and Statistical Parametric Mapping (SPM) 12 toolboxes. This process included motion and distortion correction, spatial normalization to Montreal Neurological Institute (MNI) space, and smoothing with a 6-mm full-width at half-maximum Gaussian kernel. Data quality control (QC) was then conducted on the preprocessed data to select subjects with optimal normalization to MNI space for further analysis.

### 2.3. Estimation of multi-scale ICNs

We applied a spatially constrained ICA method, MOO-ICAR [15], to each preprocessed adolescent rsfMRI dataset to extract subject-specific spatial component maps and time courses (TCs) using the NeuroMark 2.2 template (https://trendscenter.org/data/) [10]. Implementation was done using the Group ICA of fMRI Toolbox (GIFT) v4.0c (http://trendscenter.org/software/gift/) [16]. This approach leverages reference data to enhance component identification and ensure consistent ICN extraction across datasets [10], [15], [17].

### 2.4. Static Functional Network Connectivity (sFNC) Analysis

Using TCs extracted via MOO-ICAR, controlled for head motion, we computed the sFNC by estimating the Pearson correlation coefficient between each pair of ICNs’ TCs [14], yielding a 105 × 105 symmetric connectivity matrix. Each cell represents the connectivity strength between ICN pairs. To ensure accurate intrinsic connectivity representation, sFNCs were further refined by regressing out age, sex, site, and residual head motion effects.

### 2.5. Multi-variate Brain Behavior Association

We applied PCA to sFNC features from the hypothesized subdomains to reduce dimensionality. Canonical correlation analysis (CCA) was then used to identify relationships between brain connectivity and cognitive variables. Here, *X* ∈ ℝ _*n* ×*p*_ represents the PCA-reduced FNC features, and *Y* ∈ ℝ _*n*×*p*_ represents vigilance, working memory, and CBCL scores, where *n* is the number of samples, and *p* and *q* are the variable counts for each dataset. CCA finds linear transformations of Σ_*X*_, Σ_*Y*_, and Σ_*XY*_ to maximize the correlation between their canonical covariates *Xu* and *Yv* based on their joint covariance matrix Σ_*XY*_. The standard objective function can be expressed as follows:

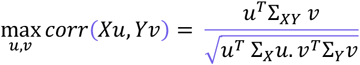

where *Xu* and *Yv* are the projections of *X* and *Y* onto the directions *u* and *v* that maximize the correlation. Pearson correlation was used to assess associations between canonical components and cognitive scores.

To ensure robust model performance, we implemented a 5-fold cross-validation approach. The data was divided into five equal subsets (“folds”), and in each iteration, the model was trained on four folds (80%) and validated on the remaining fold (20%). This process was repeated five times, with each fold serving as the validation set once. We then selected the best feature sets and identified the optimal canonical weights that maximized the association between the canonical components (CC) derived from CCA and the three cognitive and behavioral variables combined.

## 3. RESULTS

### 3.1. PCA+CCA: Multivariate brain-behavior association

We started with 5,460 sFNC features, reducing them to 630 by focusing on four subdomains. PCA further reduced dimensionality to 61 features, capturing 80% of the variance. CCA analysis on discovery sets revealed that the first canonical component (CC1) showed a strong, stable correlation with cognitive and behavioral variables, (0.228 to 0.239) shown in Fig. 1, with highly significant p-values. This association was validated in the replication set. CC2 showed moderate significance in the discovery set but had limited replication success, while CC3 exhibited high p-values across both datasets, indicating no meaningful associations.

**Fig. 1.**
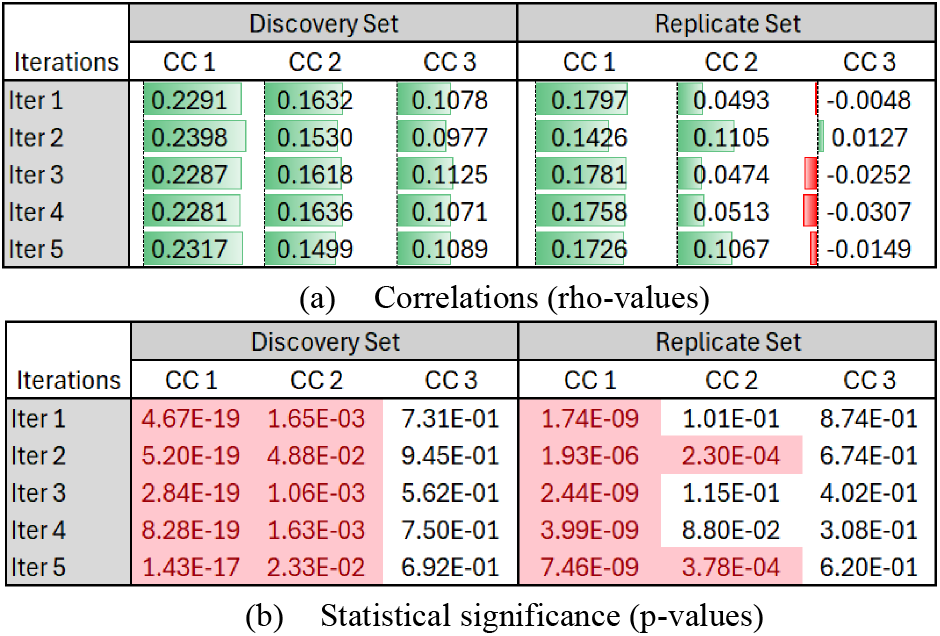
CCA analysis correlations and statistical significance. CCA analysis on PCA features and cognition outputs three canonical variables, across 5 iterations of 5-fold validation (4 folds discovery set, 1 fold replicate set). Extracted canonical variables of PCA features and cognition (a) correlation values and (b) p-values are represented above, for discovery and replicate sets.

In summary, the most substantial brain-behavior association lies within CC1 across both the discovery and replication datasets. This component captures the strongest relationship between multi-scale ICNs and cognitive-behavioral domains, marking it as the primary association pattern.

### 3.2. First Canonical Component Analysis

A closer examination of CC1 revealed that the mean loadings for vigilance, working memory, and scores were -5.28 ± 0.56, -5.24 ± 0.6, and 0.1 ± 0.01, respectively. The loading for each fold is shown in Fig. 2. The low variance in these loadings across all folds further supports the robustness of CC1 as a meaningful indicator of the association between ICNs and cognitive-behavioral measures. The scatter plots in Fig.3 illustrate the variability of CC1’s FNC feature loadings in relation to cognitive variables for both discovery and replication datasets, providing insight into the consistency of these associations. Additionally, Fig. 4 presents the loadings of all 630 features, offering a comprehensive view of the feature contributions to CC1. Finally, the connectogram in Fig. 5 highlights the top 10% of FNC feature loadings, visually representing the most influential connections within the ICN subdomains that contribute to this primary canonical component.

**Fig. 2.**
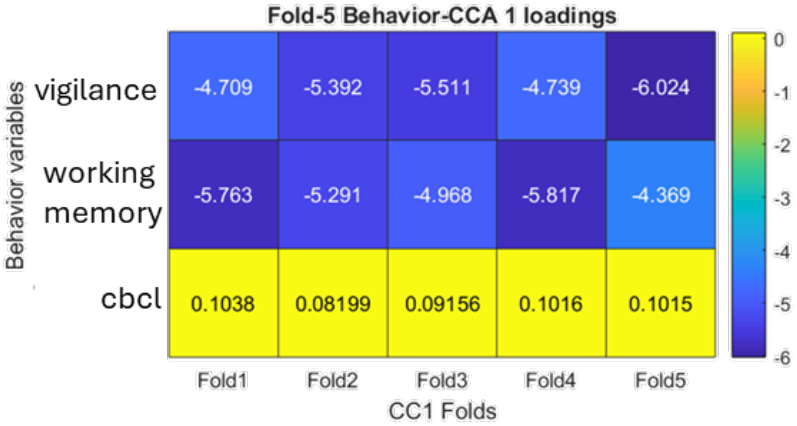
Cognition variable loadings. CCA analysis on PCA features and cognition outputs 3 canonical variables, across 5 iterations of 5-fold validation (4 folds discovery, 1 fold replicate). The figure illustrates CC1 loadings of cognition variables across all iterations in the discovery set.

**Fig. 3.**
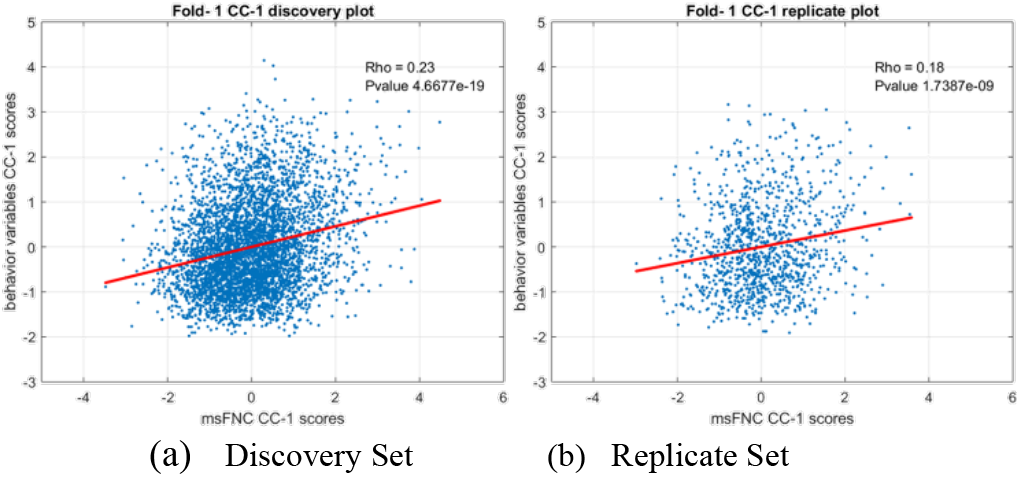
CC1 variable scores scatter plots in discovery and replicate sets. CCA analysis on PCA features and cognition outputs three canonical variables, across 5 iterations of 5-fold validation (4 folds discovery, 1 fold replicate). (a) Iteration 1, Scatter plot of CC1 scores for FNC PCA features and cognition variables in the discovery set. (b) Iteration 1, Scatter plot of CC1 scores for FNC PCA features and cognition variables in the replication set. The X-axis represents FNC PCA feature scores, and the Y-axis represents cognition variable scores.

**Fig. 4.**
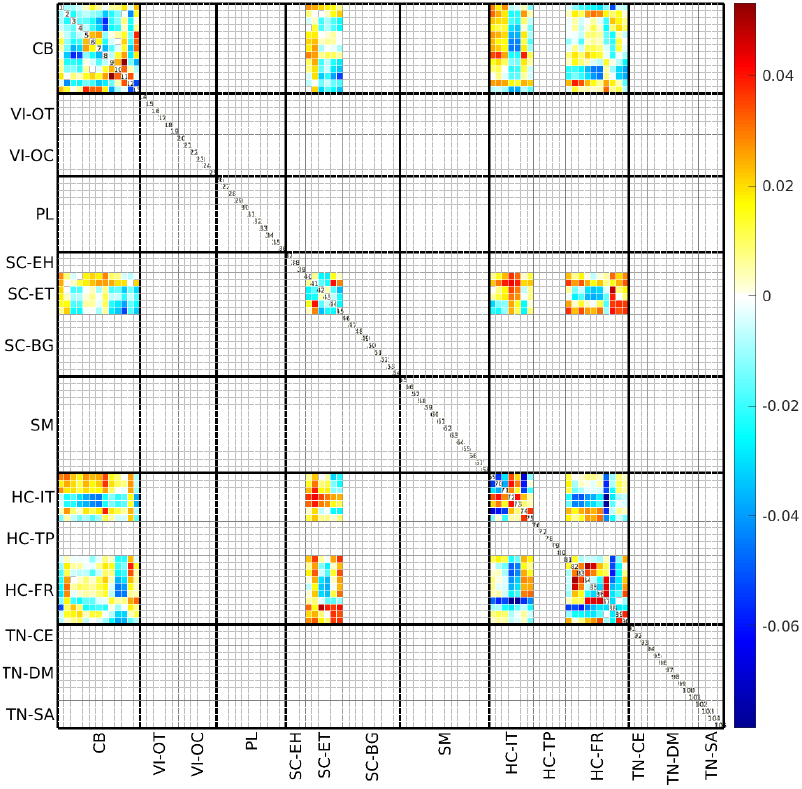
FNC feature loadings. CCA analysis on PCA features and cognition outputs 3 canonical variables, across 5 iterations of 5-fold validation (4 folds discovery, 1 fold replicate). The figure illustrates iteration 1, where CC1 loadings are mapped from PC space to FNC space and highlights connectivity between multi-scale ICNs across CB, SC-ET, HC-IT, and HC-FR subdomains

**Fig. 5.**
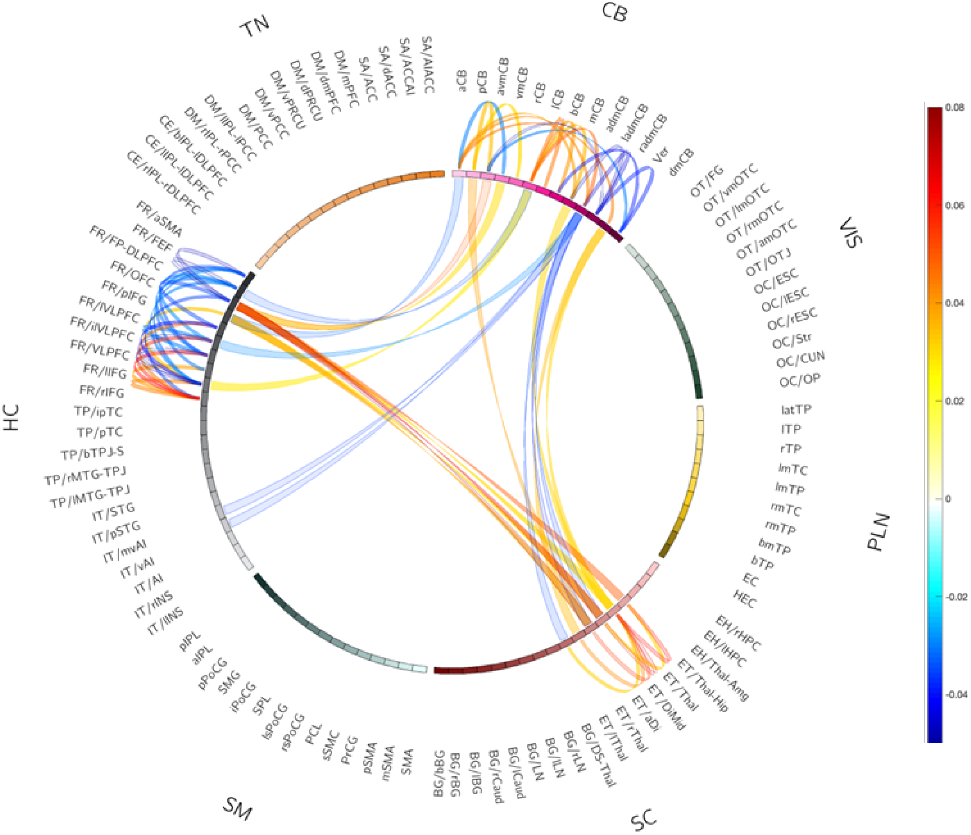
FNC features loadings. CCA analysis on PCA features and cognition outputs three canonical variables, across 5 iterations of 5-fold validation (4 folds discovery set, 1 fold replicate set). The figure represents iteration 1, where CC1 loadings of FNC PCA features are mapped back to FNC space. The connectogram visualizes the top 10% of CC loadings, highlighting key ICN connections.

These findings suggest that CC1 not only captures the strongest association between multi-scale ICNs and cognitive-behavioral domains but also highlights specific connectivity patterns that may underlie these relationships, offering potential insights into brain-based markers of cognitive and behavioral functions.

## 4. DISCUSSION

This study examines the relationship between sFNC from multi-scale ICNs and cognitive-behavioral domains in adolescents. Our findings highlight significant associations between connectivity patterns and cognitive functions, particularly vigilance and working memory

The top 10% of loadings highlight substantial intra-subdomain connections, with HC-FR exhibiting 20 ICNs, CB 16 ICNs, SC-ET 6 ICNs, and HC-IT none. For inter-subdomain connectivity, CB and SC-ET had 7 connections, with CB and HC-FR, and HC-FR and SC-ET pairs each had 6. In contrast, HC-IT had only 2 connections with CB. This disparity suggests that specific ICN configurations are pivotal for cognitive functions, underscoring the importance of intra-subdomain connectivity.

Furthermore, our findings emphasize the effectiveness of combining PCA and CCA for dimensionality reduction and feature extraction in high-dimensional neuroimaging data. This integrated approach not only simplifies data complexity but also preserves meaningful variability in the relationships between sFNC and cognitive outcomes. By providing a clearer understanding of the neural contributions to cognitive and behavioral functions, our findings support a multi-scale perspective of brain organization during adolescence, which has implications for identifying potential markers of neurodevelopmental disorders.

## 5. CONCLUSION

This study highlights the role of multi-scale ICNs in cognitive and behavioral profiles of adolescents, with the first canonical component showing the strongest associations with attention, working memory, and behavioral regulation. Consistent findings across discovery and replication sets support the stability of these relationships, emphasizing the potential of multi-scale connectivity analysis in understanding neurodevelopmental disorders like ADHD. Future research should explore mechanisms and clinical relevance, particularly for assessing neurodevelopmental risk related to attentional and memory deficits.

## 6. COMPLIANCE WITH ETHICAL STANDARDS

Informed written consent was obtained from children and their parents, with ethical approval from each research site’s institutional review boards.

## 7. ACKNOWLEDGMENTS

This work was supported by grants NSF (2316421 and 2112455), awarded to Dr. Vince Calhoun, NIH R01MH130595, awarded to Dr. Jingyu Liu, and grant NIH R01MH119251, awarded to Dr. Armin Iraji.

